# Evaluating non-oxidative glycolysis using ^13^C metabolic flux analysis

**DOI:** 10.64898/2026.11.04.764406

**Authors:** Kenta Miyoshi, Tatsumi Imada, Haruka Teraki, Teppei Niide, Yoshihiro Toya, Hiroshi Shimizu

## Abstract

Non-oxidative glycolysis (NOG) converts glucose to acetyl-CoA without emitting CO_2_ during *Escherichia coli* bioproduction. Although ^13^C-metabolic flux analysis (^13^C-MFA) has the potential to evaluate NOG flux distribution *in vivo*, the precision of these estimates depends on the labeling patterns of glucose tracers. Because *E. coli* has multiple native glycolysis pathways, selecting the optimal glucose tracer is important for precise estimates. This study used simulations to evaluate the performance of 18 commercially available glucose tracers in precisely predicting NOG flux. We artificially generated flux distributions of central metabolic pathways, including three types of NOG pathways, and obtained ^13^C labeling patterns of amino acids metabolized from each glucose tracer. Using these data, we then calculated 95% confidence intervals (CIs) for NOG pathway fluxes. Comparing the sum of CI widths from three NOG fluxes showed that estimation with [1,3-^13^C]-glucose yielded the narrowest CIs. Several tracers, including [2,3-^13^C]glucose, were capable of determining fluxes with high precision, but [1-^13^C]glucose was not one of them, despite often being used for estimating glycolytic fluxes. We then experimentally evaluated [2,3-^13^C]- and [1-^13^C]glucose as tracers in *E. coli* strains with and without the expression of phosphoketolase, which is required for NOG pathways to function. The results suggested that [1-^13^C]glucose could not detect NOG fluxes, whereas [2,3-^13^C]glucose estimated the fluxes of all three NOG pathways with narrow CIs.

## 1 Introduction

Microorganisms have beneficial industrial applications through their ability to enhance the carbon yield of a desired product in biorefineries. Acetyl-CoA is an important precursor involved in the synthesis of various valuable products. Hence, increasing acetyl-CoA supply is a promising strategy for metabolic engineering. Most organisms, including *Escherichia coli*, possess multiple glycolytic pathways that catabolize glucose into acetyl-CoA, such as the Embden-Meyerhof (EM), pentose phosphate (PP), and Entner-Doudoroff (ED) pathways. However, these pathways proceed via a pyruvate dehydrogenase reaction that loses carbon through CO_2_ release, a limiting factor in the production of acetyl-CoA derivatives (1). To avoid carbon loss, an alternative route for converting glucose to acetyl-CoA is non-oxidative glycolysis (NOG). These pathways appear to act through converting one glucose molecule into three acetyl-CoA molecules. *Bifidobacterium*-derived phosphoketolase has been employed to form NOG (Figure 1) (2,3).

**Figure 1.**
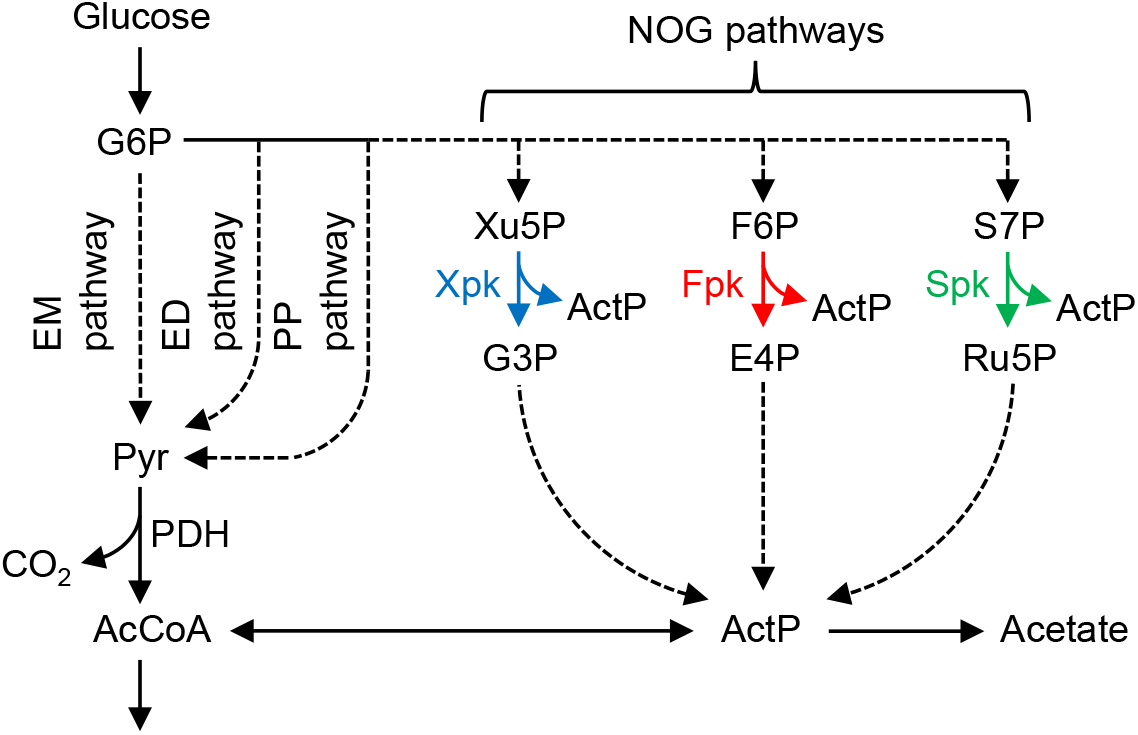
Overview of *E. coli* glycolytic pathways, including three variations of NOG pathways. This study simulated reactions with 18 commercially available glucose tracers to evaluate the precision of estimating NOG pathway flux. Using a model that incorporates atom transitions for each reaction, we generated a virtual dataset comprising mass isotopomer distributions (MIDs) of amino acids from the hypothetical flux distributions. Flux estimation was performed using this virtual MID dataset and 95% confidence intervals (CIs) of NOG pathway fluxes per glucose tracer were compared. From these results, promising glucose tracers were identified and used in cultures of *E. coli* expressing phosphoketolase; these experiments aimed to verify that NOG pathway flux could be detected.

Once NOG pathways are induced in a host cell, their *in vivo* reaction rates should be quantified to advance cell engineering. ^13^C-metabolic flux analysis (^13^C-MFA) can experimentally evaluate the flux distribution of metabolic pathways based on ^13^C-labeling experiments (4-6). In this method, cells are grown on substrates with specific ^13^C-labeled sites. Because the labeled substrate is metabolized through various pathways, flux distribution can be estimated from the resultant metabolite labeling patterns. A precise (i.e., narrow confidence interval) estimate of a reaction’s flux requires a substrate with ^13^C at an appropriate carbon position (7). [1-^13^C]glucose is widely used to distinguish multiple glycolytic pathways including the EM, PP, and ED pathways (8). However, as the complexity of the glucose catabolic routes increases when considering the NOG pathways, the appropriate glucose tracer for ^13^C-MFA is unclear. A previous study used [1-^13^C]glucose to estimate flux in a NOG-expressing *E. coli* strain (9), but it is uncertain whether this tracer is the appropriate choice for evaluating the flux of all NOG pathways. In cyanobacteria, [1,2-^13^C]glucose is sometimes used because they possess the Calvin-Benson cycle in addition to the EM and PP pathways (10). It has also been reported that evaluating phosphoketolase flux in cyanobacteria requires parallel labeling experiments using four types of glucose tracers (11). Furthermore, phosphoketolase is known for its promiscuity, catalyzing not only fructose 6-phosphate (F6P) and xylulose 5-phosphate (X5P) but also sedoheptulose 7-phosphate (S7P) (12). Consequently, when analyzing phosphoketolase-expressing *E. coli*, the selection of an appropriate tracer requires considering multiple complex glycolytic pathways (e.g., EM, PP, ED) and three variations of NOG pathways (Figure 1).

### Materials and methods Strains and culture conditions

Strain MKEC03, engineered to activate NOG pathways, and its control strain MKEC01 were used (13). MKEC03 possesses a plasmid for expressing two enzymes under the T7 promoter: a codon-optimized *Bifidobacterium adolescentis*-derived phosphoketolase (*fxpk*) and *E. coli*-derived fructose bisphosphatase (*fbp*). MKEC01 contains an empty vector. Both strains were inoculated from glycerol stock into 5 mL LB medium (10 g L^−1^ tryptone, 5 g L^−1^ yeast extract, and 5 g L^−1^ NaCl) and cultured aerobically in test tubes (37°C, 150 rpm). Overnight cultures were transferred to fresh 50 mL M9W medium (13) in a 200-mL baffled Erlenmeyer flask with an initial optical density (OD_600_) of 0.05 at 600 nm, and grown aerobically (37°C, 150 rpm; BF-43FL; TAITEC, Saitama, Japan). All cultures were supplemented with 0.1 mM IPTG as the inducer and 0.05 g L^−1^ ampicillin.

### Cell growth and extracellular metabolites

Cell concentrations in the culture broth were measured at OD_600_ using a UV-mini 1240 spectrophotometer (Shimadzu, Kyoto, Japan). The culture broth was centrifuged (4°C, 1min, 17,800×*g*) and passed through a Millex-HV 0.45 μm filter (Merck Millipore, Billerica, MA, USA) to obtain the supernatant. The culture broth’s acetate concentration was measured using an Aminex HPX-87H column (Bio-Rad, Hercules, CA, USA) in a high-performance liquid chromatograph (Shimadzu) equipped with a refractive index detector (RID-20A) and UV/Vis detector (SPD-20A). The procedure followed published methods (8). Glucose concentrations were measured in an enzyme electrode sensor BF-5 (Oji Scientific Instruments, Hyogo, Japan).

### Gas chromatography-mass spectrometry of proteinogenic amino acids

Cell pellets were obtained from cultures through centrifugation (4°C, 5 min, 4,000×*g*) to achieve an OD_600_ × volume (mL) ≒ 3, followed by hydrolysis using 6N HCl (105°C, 18 h). Proteinogenic amino acids were converted to their *tert*-butyldimethyl derivatives and analyzed with gas chromatography-mass spectrometry (GC-MS-QP2020 NX, Shimadzu) using a DB-5MS+DG column (Agilent Technologies). The procedure followed published methods (14).

### ^13^C-Metabolic flux analysis

The metabolic pathways used in ^13^C-MFA included the EM, PP, and ED pathways, TCA cycle, glyoxylate shunt, anaplerotic reactions, and three types of NOG reactions (Xpk, Fpk, and Spk). Reaction stoichiometries and their carbon-atom transitions are listed in Table S1. Reactions and carbon-atom transitions from precursor metabolites to amino acids are listed in Table S2. The elementary metabolite unit framework was used to develop a model for calculating MIDs from flux distributions (15). Precursor requirements for biomass formation were obtained from the literature (16). Flux distributions were estimated such that the residual sum of squares (RSS) is minimized between measured and simulated amino acid MIDs. Equation 1 was used as the function for ^13^C-MFA to evaluate the flux estimation precision for each glucose tracer. Equation 2 was used to determine efflux perturbations in ^13^C-MFA.

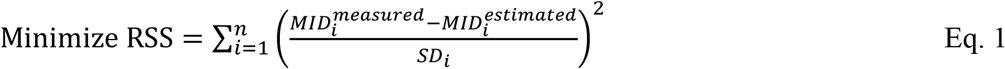

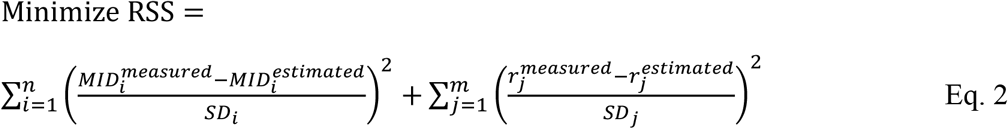

where MID_i_^measured^ and MID_i_^estimated^ represent the measured and simulated MID of *i*th amino acid fragment, respectively. The standard deviation (SD) of the MIDs was set to 0.01 for all fragments (8). In Equation 2, r_j_^measured^ and r_j_^estimated^ represent measured and simulated effluxes of the *j*th reaction, respectively. Efflux SD was calculated from replicate measurements. The goodness of fit between measured and simulated MIDs was evaluated using *χ*^2^ tests. The 95% CIs of estimated fluxes were determined using a grid-search algorithm (17). To evaluate ^13^C-tracers in simulations, amino acid MIDs were generated from three types of assumed flux distributions in OpenMebius (18). ^13^C-MFA was performed in OpenMebius2 (version 3.5.0) (19).

## Results and Discussion

### Growth profiles of *E. coli* strains with and without NOG

We cultured MKEC03 in a synthetic medium with glucose as the sole carbon source using the MKEC03, which expressed *fxpk* and *fbp* to function NOG, and the control MKEC01. Figure 2 depicts the time courses of OD_600_, glucose, and acetate concentrations. The specific rates of growth, glucose consumption, and acetate formation are summarized in Table 1. MKEC03 had lower specific rates than MKEC01, but culture profiles alone were insufficient to determine whether NOG pathways functioned *in vivo*.

**Table 1.** Specific rates of MKEC01 and MKEC03 strains.

| Strain | Cell growth<br>(h<sup>-1</sup>) | Glucose consumption<br>(mmol g<sup>-1</sup> h<sup>-1</sup>) | Acetate production<br>(mmol g<sup>-1</sup> h<sup>-1</sup>) |
|---|---|---|---|
| MKEC01 | 0.68 ± 0.02 | 8.6 ± 0.3 | 3.7 ± 0.4 |
| MKEC03 | 0.56 ± 0.01 | 7.7 ± 0.3 | 3.3 ± 0.1 |

**Figure 2.**
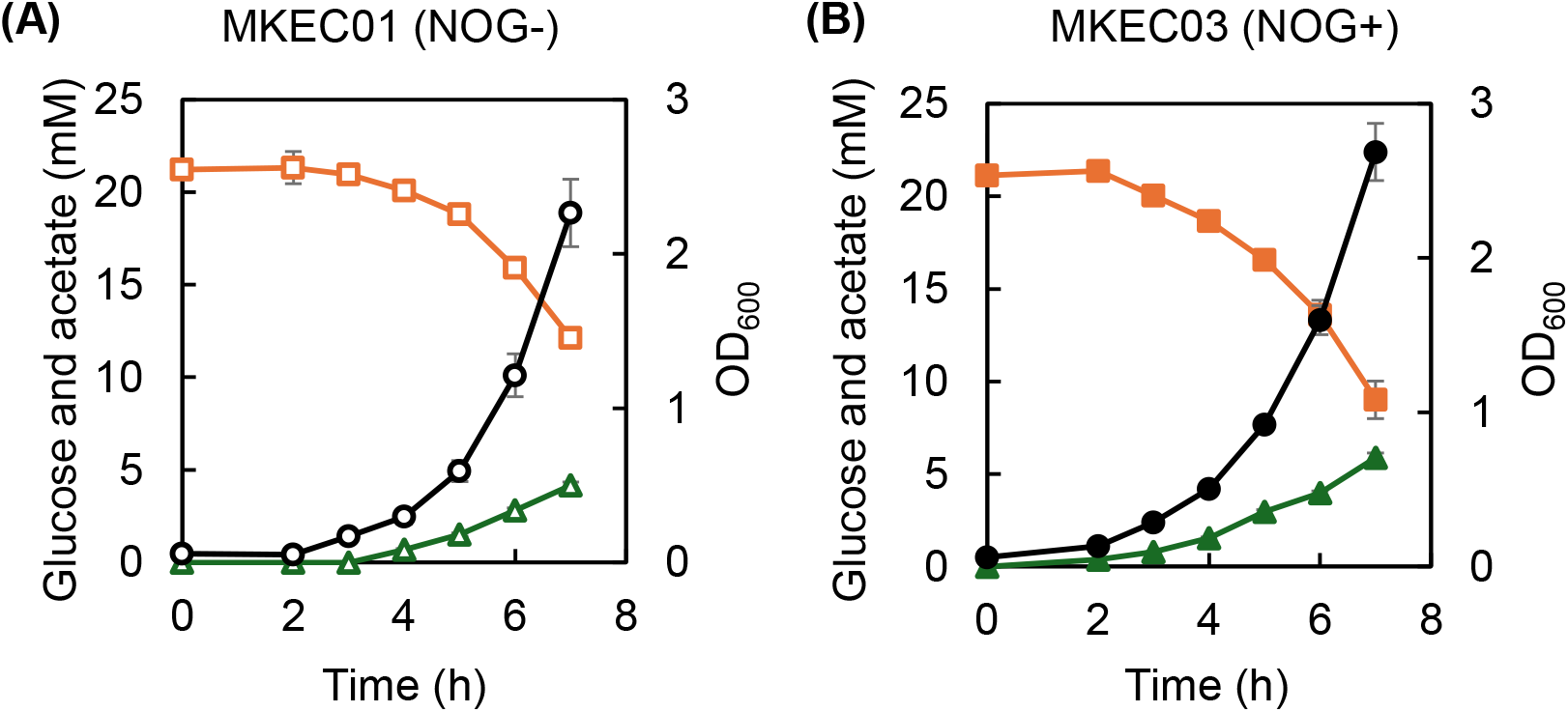
Batch-culture profiles of (A) MKEC01 and (B) MKEC03 strains. Open, square, and triangular symbols represent the OD_600_, glucose concentration, and acetate concentration, respectively.

### Comparison of ^13^C-glucose labeling for evaluating flux of the NOG pathways

The precision of ^13^C-MFA flux estimations is represented by the 95% CI widths, and is affected by which carbon atom of glucose is labeled, the pathway structure, and flux distribution to be estimated. Although [1-^13^C]glucose is often used to distinguish carbon derived from the EM, PP, and ED pathways, the additional network complexity of including NOG pathways means we cannot assume that [1-^13^C]glucose remains the best tracer for high-precision estimation.

We tested 18 commercially available glucose tracers (Table S3) using simulations to determine the precision of their NOG flux predictions. A pathway model involving the Xpk, Fpk, and Spk reactions was developed. Additionally, a flux distribution was required to conduct virtual experiments. Therefore, using the specific growth rate of MKEC03 (Table 1), we generated three types of flux distributions: high Xpk, high Fpk, and high Spk (Figure S1). For each of these flux distributions, we performed a virtual ^13^C-labeling experiment using the 18 glucose tracers and generated amino acid MIDs (Table S4). ^13^C-MFA was performed using the virtual MID dataset, and 95% CIs of the Xpk, Fpk, and Spk fluxes were determined.

Figure 3 shows the 95% CIs widths for Fpk, Xpk, and Spk fluxes per glucose tracer, across three hypothetical flux distributions. The feasible flux range for each NOG pathway was 10.75 mmol g^-1^ h^-1^, suggesting that [4-^13^C]glucose did not narrow the CI of NOG pathway fluxes. Similarly, [4-^13^C]glucose was the least suitable labeled substrate in a prior experiment investigating the optimal tracer for ^13^C-MFA in *E. coli* without considering NOG pathways (20). Hence, [4-^13^C]glucose appears unsuitable with or without NOG pathways. The narrowest CIs occurred with [1,3-^13^C]glucose for all three flux distributions, followed by [2,3-^13^C]-, [5,6-^13^C]-, [1,2,3-^13^C]-, and [4,5,6-^13^C]glucose. The widely used [1,2-^13^C]glucose was still capable of determining flux, whereas [1-^13^C]glucose yielded broader CIs that indicated it was unsuitable for detection.

**Figure 3.**
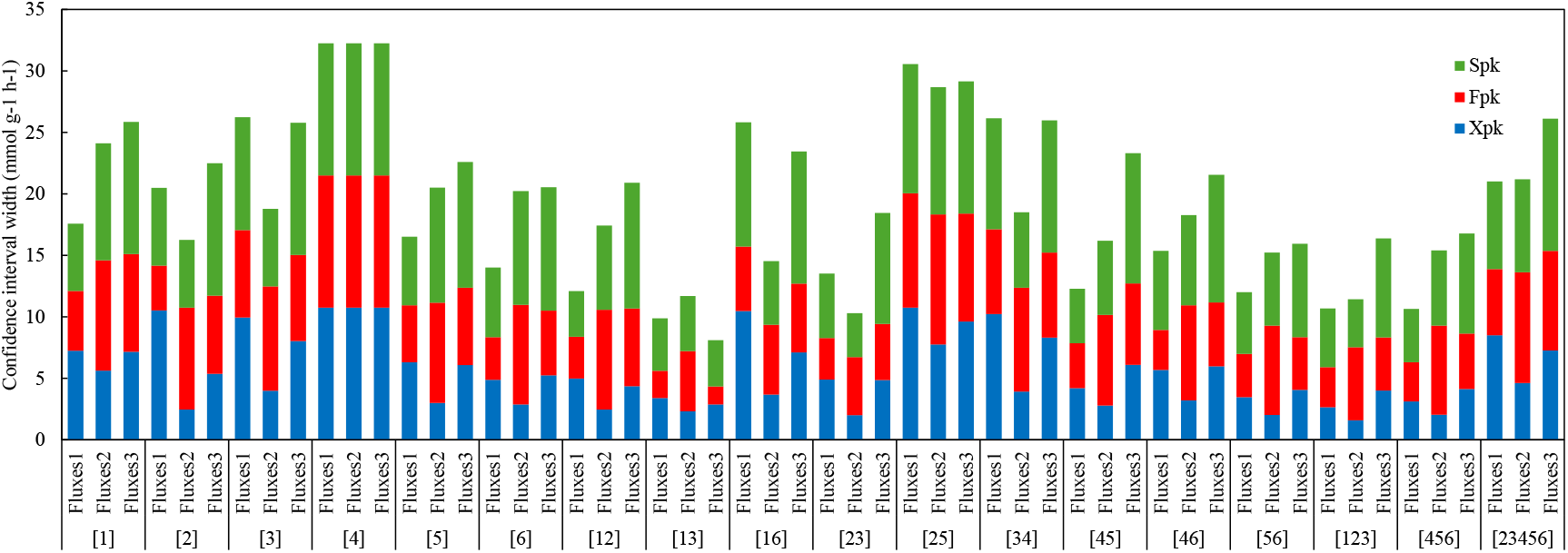
95% Confidence-interval widths of NOG pathways fluxes with 18 different glucose tracers from *in silico* ^13^C-labeling experiments. Blue, red, and green bars represent CI widths of Xpk, Fpk, and Spk fluxes, respectively. Labels indicate the ^13^C positions in glucose (e.g., [1], [1-^13^C]glucose; [12], [1,2-^13^C]glucose). On the horizontal axis, fluxes 1, 2, and 3 indicate artificial distributions with high Xpk, Fpk, and Spk fluxes, respectively.

### Experimental verifications of ^13^C-MFA using phosphoketolase-expressing *E. coli*

*In silico* simulations indicated that certain tracers (e.g., [1,3-^13^C]- and [2,3-^13^C]glucose) enabled the estimation of Fpk, Xpk, and Spk fluxes with narrow CIs, whereas [1-^13^C]glucose did not. We then selected [1-^13^C]-, [1,2-^13^C]-, and [2,3-^13^C]glucose for the experimental verification of NOG flux estimations, as these three tracers were readily available in our laboratory stock. We separately cultured MKEC01 and MKEC03 cells with each labeled glucose and analyzed their proteinogenic amino acids using GC/MS. Table S5 shows amino acid MIDs corrected for natural isotope abundance, compared across measured and simulated values. We then applied *χ*^2^ tests to assess the fit between measured and estimated MIDs. The threshold for this goodness-of-fit *χ*^2^ test was 97. The RSS values from using [1-^13^C]-, [1,2-^13^C]-, and [2,3-^13^C]glucose were 16, 27, and 26 for MKEC01, respectively, as well as 21, 70, and 119 for MKEC03. For MKEC03 with [2,3-^13^C]glucose, RSS slightly exceeded the threshold, and the fit did not pass the *χ*^2^ test. Therefore, the results for this condition are presented as reference values. Nevertheless, the trend in CI width is noteworthy.

Figure 4 illustrates the 95% CIs for NOG fluxes corresponding to each glucose tracer. Consistent with prior simulations, CI widths diminished sequentially from [1-^13^C]-to [1,2-^13^C]- and finally to [2,3-^13^C]glucose. In particular, when [1-^13^C] glucose was used with MKEC03, the 95% CI for Xpk, Fpk, and Spk fluxes included zero, meaning NOG activity could not be ascertained. In contrast, when [1,2-^13^C]- or [2,3-^13^C]glucose was used, a flux greater than zero was detected through the Fpk reaction, suggesting that the NOG pathway was functional. In MKEC01, the CIs for Xpk, Fpk, and Spk calculated using [1-^13^C]glucose were wide. When [2,3-^13^C]glucose was used, none of the estimated fluxes had narrow CIs.

**Figure 4.**
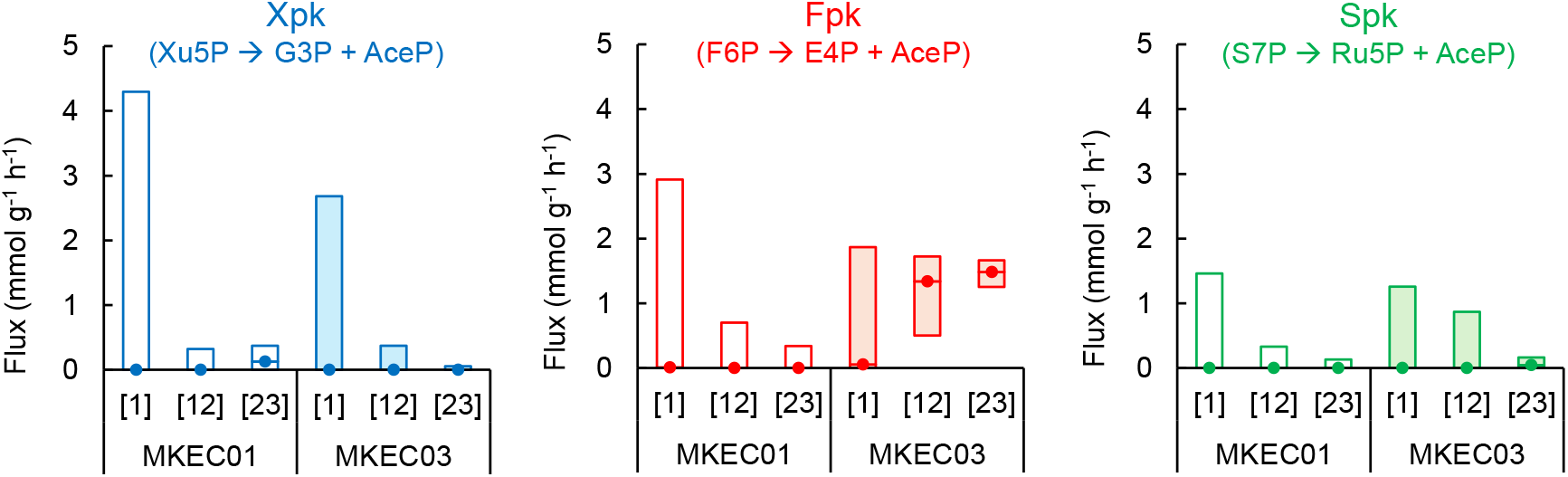
Confidence intervals of Xpk, Fpk, and Spk fluxes in MKEC01 and MKEC03. Boxes represent the 95% CIs of estimated fluxes. The points indicate best-fit flux values. The labels indicate the positions of ^13^C in glucose (e.g., [1], [1-^13^C]glucose; [12], [1,2-^13^C]glucose).

^13^C-MFA revealed that only the Fpk reaction was active in MKEC03. The Michaelis constants (Km) of *B. adolescentis*-derived phosphoketolase for X5P, F6P, and S7P are 6.7, 22.7, and 68.1 mM, respectively (12). Given the high Km value for S7P, it was not surprising that the Spk flux was low. Although the lower Km for X5P suggests that the Xpk reaction with X5P occurred more readily, the Fpk reaction with F6P proceeded preferentially, probably because Fbp expression enhanced F6P supply. The total concentration of pentose 5-phosphates (X5P, Ru5P, and R5P) in *E. coli* is approximately 1.3 mM (21). Hence, low intracellular X5P concentrations may explain the limited contribution of Xpk.

Figure 5 shows overall flux distributions of MKEC01 and MKEC03 using the [1,2-^13^C]glucose dataset. Although Fpk flux was observed in MKEC03, its value was small (range = 0.5–1.7 mmol g^-1^ h^-1^) compared with phosphofructokinase (Pfk) flux (range = 4.4–5.2 mmol g^-1^ h^-1^) of the EM pathway, which competes for F6P utilization. To increase Fpk flux, Pfk flux must be suppressed. In a previous study, enhanced acetate yield in phosphoketolase-expressing *E. coli* was achieved through deleting *pfkA* (encodes Pfk), suggesting an increased flux through the Fpk reaction (13). Hence, to activate NOG pathways, the accumulation of their precursors must be triggered through engineering.

**Figure 5.**
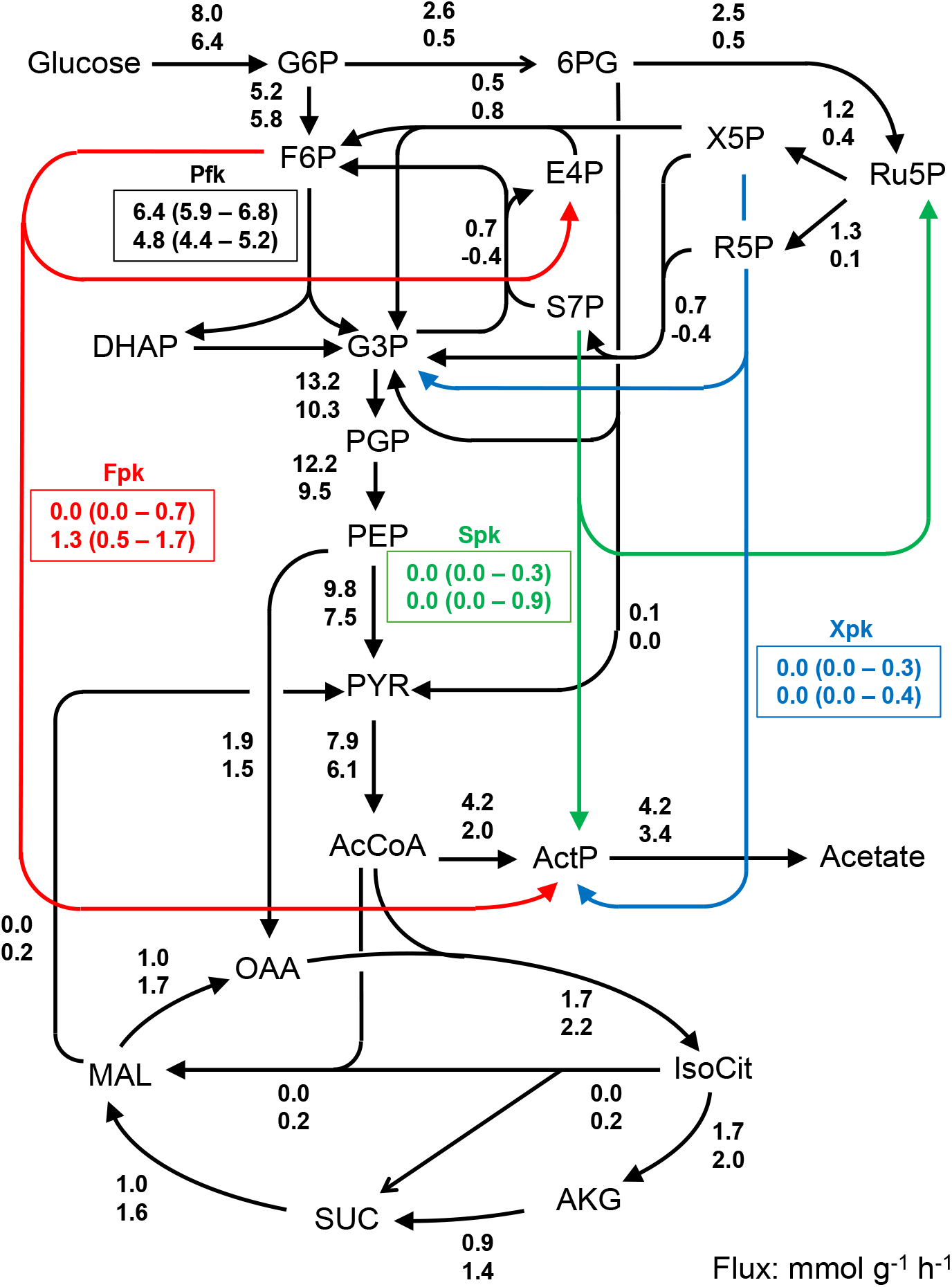
Flux distributions of MKEC01 and MKEC03 by [1,2-^13^C] glucose. Upper and lower values represent fluxes (mmol g^-1^ h^-1^) of MKEC01 and MKEC03, respectively. Values in parentheses indicate 95% CIs. Metabolite abbreviations: 6PG, 6-phosphogluconate; AcCoA, acetyl-CoA; ActP, acetyl phosphate; AKG, 2-ketoglutarate; DHAP, dihydroxyacetone phosphate; E4P, erythrose 4-phosphate; F6P, fructose 6-phosphate; G6P, glucose 6-phosphate; G3P, glyceraldehyde 3-phosphate; IsoCit, isocitrate; MAL, malate; OAA, oxaloacetate; PEP, phosphoenolpyruvate; PGP, 3-phosphoglycerate; PYR, pyruvate; R5P, ribose 5-phosphate; Ru5P, ribulose 5-phosphate; S7P, sedoheptulose 7-phosphate; SUC, succinate; X5P, xylulose 5-phosphate.

In this study, we predicted ^13^C glucose labeling patterns that enable high-precision estimation of NOG fluxes and experimentally validated the flux-detection ability of those predicted ^13^C tracers in phosphoketolase-expressing *E. coli*. These techniques and findings will benefit efforts to perform metabolic engineering aimed at exploiting the NOG pathways in *E. coli*.

## Acknowledgments

This research was supported by Program for Leading Graduate Schools (Humanware Innovation Program, Osaka University, Japan); a Grant-in-Aid for Scientific Research (S) (No. 19H05626; No. JP25K24646); Scientific Research (B) No. 25K01591; Scientific Research (C) No. 25K08399 from the Japan Society for the Promotion of Science (JSPS); Green Technologies of Excellence (GteX) Program Japan Grant Number JPMJGX23B4; Adopting Sustainable Partnerships for Innovative Research Ecosystem (ASPIRE) Grant Number JPMJAP24A2 from the Japan Science and Technology Agency (JST); and a research grant from the Institute for Fermentation, Osaka (IFO).

## Author contributions

K.M.: Investigation, Formal analysis, Data curation, Writing – original draft. T.I.: Data curation, Software, Validation, Writing – review & editing. H.T.: Formal analysis, Writing – review & editing. T.N.: Validation, Writing – review & editing. Y.T.: Conceptualization, Formal analysis, Methodology, Project administration, Visualization, Writing – original draft. H.S.: Conceptualization, Funding acquisition, Supervision, Writing – review & editing.

## Declarations

Conflict of interest: The authors declare no competing interests.

Ethical approval: Not applicable. This study did not involve humans or animals.

## Data availability

Data supporting this study are included with in the manuscript or supplementary information file.

