## Supplementary material for "Evaluating non-oxidative glycolysis using ^13^C metabolic flux analysis": Fig. S1

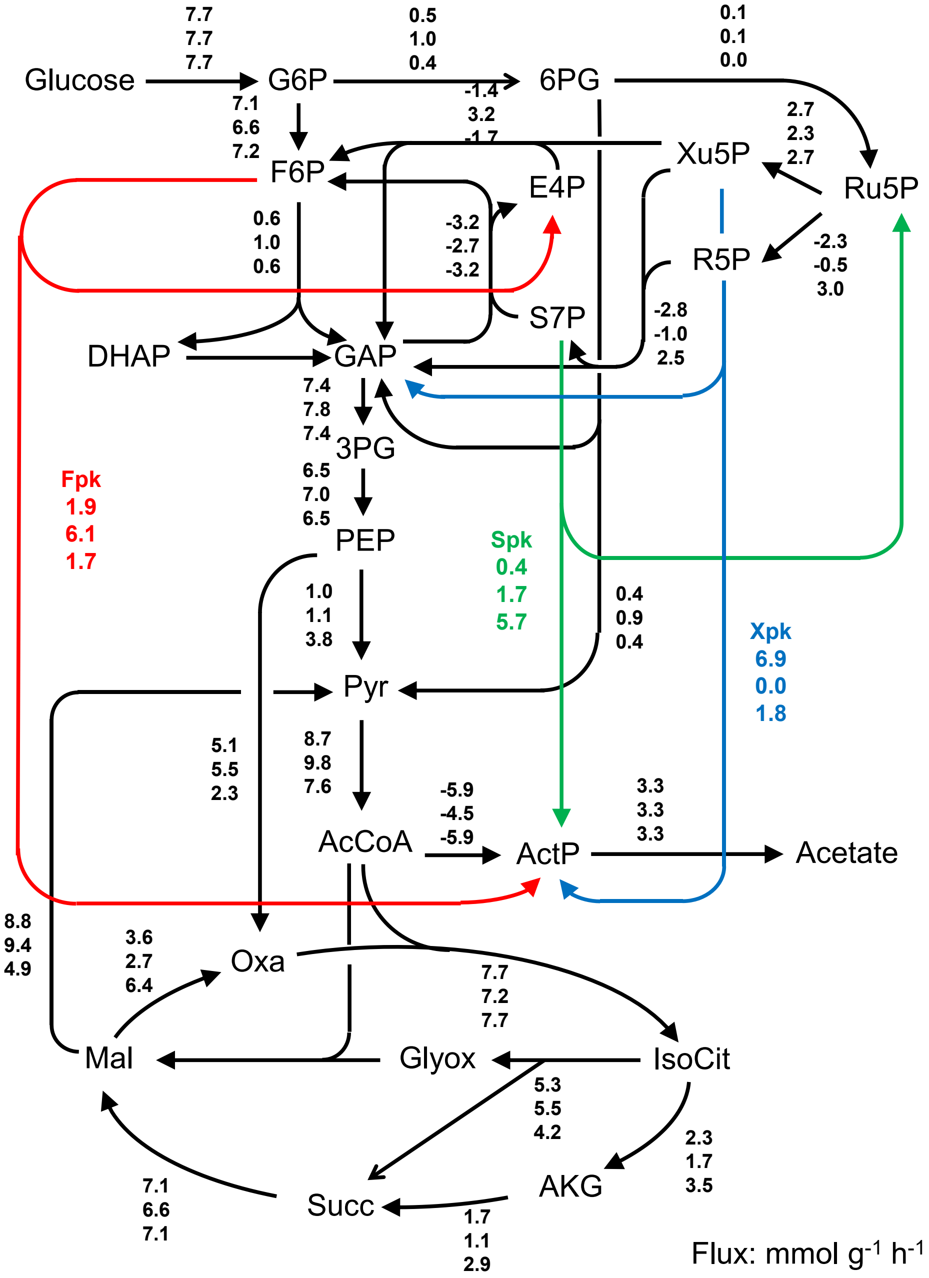


**Figure S1 Hypothetical three types of flux distributions for simulations.**
Top, middle, and bottom values represent fluxes (mmol $g^{-1}$ $h^{-1}$) of Fluxes 1 (high Xpk), Fluxes 2 (high Fpk), and Fluxes 3 (high Spk), respectively. Metabolite abbreviations are as in Fig. 5.
